# Virtual reality empowered deep learning analysis of brain activity

**DOI:** 10.1101/2023.05.18.540970

**Authors:** Doris Kaltenecker, Rami Al-Maskari, Moritz Negwer, Luciano Hoeher, Florian Kofler, Shan Zhao, Mihail Todorov, Johannes Christian Paetzold, Benedikt Wiestler, Julia Geppert, Pauline Morigny, Maria Rohm, Bjoern H. Menze, Stephan Herzig, Mauricio Berriel Diaz, Ali Ertürk

## Abstract

Tissue clearing and fluorescent microscopy are powerful tools for unbiased organ-scale protein expression studies. Critical for interpreting expression patterns of large imaged volumes are reliable quantification methods. Here, we present DELiVR a deep learning pipeline that uses virtual reality (**VR**)-generated training data to train deep neural networks, and quantify c-Fos as marker for neuronal activity in cleared mouse brains and map its expression at cellular resolution. VR annotation significantly accelerated the speed of generating training data compared to conventional 2D slice based annotation. DELiVR detects cells with much higher precision than current threshold-based pipelines, and provides an extensive toolbox for data visualization, inspection and comparison. We applied DELiVR to profile cancer-related mouse brain activity, and discovered a novel activation pattern that distinguishes between weight-stable cancer and cancer-associated weight loss. Thus, DELiVR provides a robust mouse brain analysis pipeline at cellular scale that can be used to study brain activity patterns in health and disease.

The DELiVR software, Fiji plugin and documentation can be found at **https://www.DISCOtechnologies.org/DELiVR/**.

**Graphical Abstract:** 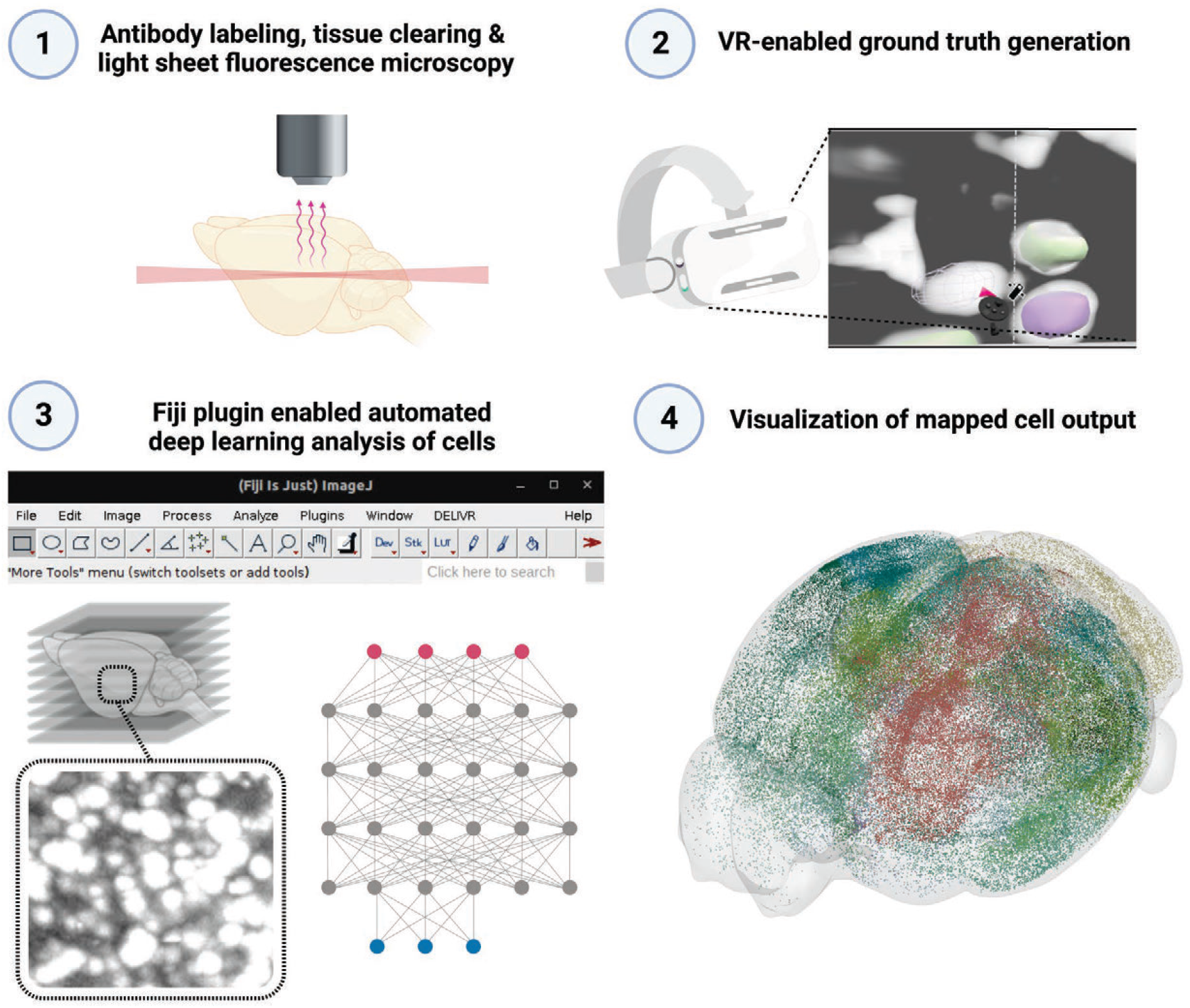

**Highlights:**

1. **DELiVR detects labelled cells in cleared brains with deep learning**
2. **DELiVR is trained by annotating ground-truth data in virtual reality (VR)**
3. **DELiVR is launched via a FIJI plugin anywhere from PCs to clusters**
4. **Using DELiVR, we found new brain activity patterns in weight-stable vs. cachectic cancer**

**Supplementary Videos can be seen at: https://www.DISCOtechnologies.org/DELiVR/**

## INTRODUCTION

Analyzing the expression of proteins is essential to understand cellular and molecular processes in physiological and disease conditions. While standard immunohistochemistry is useful to validate protein expression on tissue sections, it does not provide a holistic view of expression patterns in larger tissue pieces or whole organs. In addition, essential information can be lost during slicing^1,2^. Tissue clearing and fluorescent imaging solve many of these restrictions and allow unbiased protein expression analysis at up to organism-scale^1,3,4^. By immunostaining for the expression of immediate early genes such as c-Fos, it is possible to retrieve a brain-wide snapshot of the neuronal activity of an animal shortly before fixation. Unbiased quantification methods for system-level examination at single-cell resolution are essential to interpret those brain-wide findings^5^. Current automated methods for cell detection and registration to the Allen Mouse Brain Atlas were shown to be a valuable tool when mapping brain activity following drug treatment, whisker-evoked sensory processing, nesting or fasting^6-8^. However, these methods are commonly challenged by some aspects of 3D whole-brain imaging, specifically by variations in image acquisitions among samples, uneven signal to noise ratio across the tissue depth, or low abundance of the target protein. In such cases, applying a single threshold to a whole-brain scan can lead to a significant lack of detection sensitivity and/or specificity. As a result, current pipelines tend to use conservative thresholds and discard a lot of potentially useful information.

In response to these challenges, we developed DELiVR (**D**eep **L**earning and **Vi**rtual **R**eality mesoscale annotation pipeline), a VR aided deep learning algorithm for detecting c-Fos+ cells in cleared mouse brains (**Fig. 1**). We used the SHANEL protocol for c-Fos immunostaining^9^, tissue clearing and light-sheet fluorescence microscopy (**LSFM**). In order to analyze the resulting 3D images, we developed DELiVR: first, we generated high-quality ground-truth data by segmenting c-Fos+ cells in VR. Next, we trained a deep neural network on these ground truth data, and used it to identify cells across the brain. Subsequently, we registered the image stacks to the Allen Brain Atlas in reference-brain space, and developed a pipeline to map the aligned cells back into the original image space. We used DELiVR to map cancer-related brain activity in tumor-bearing mice that were either weight-stable or displayed cancer-associated weight loss (cachexia). Interestingly, DELiVR revealed increased neuronal activity in mice with weight-stable cancer in brain areas related to sensory processing and foraging. In contrast, this increase was lost in animals with weight loss, suggesting a weight-stable cancer-specific neurophysiological hyper-activation phenotype.

**Figure 1:**
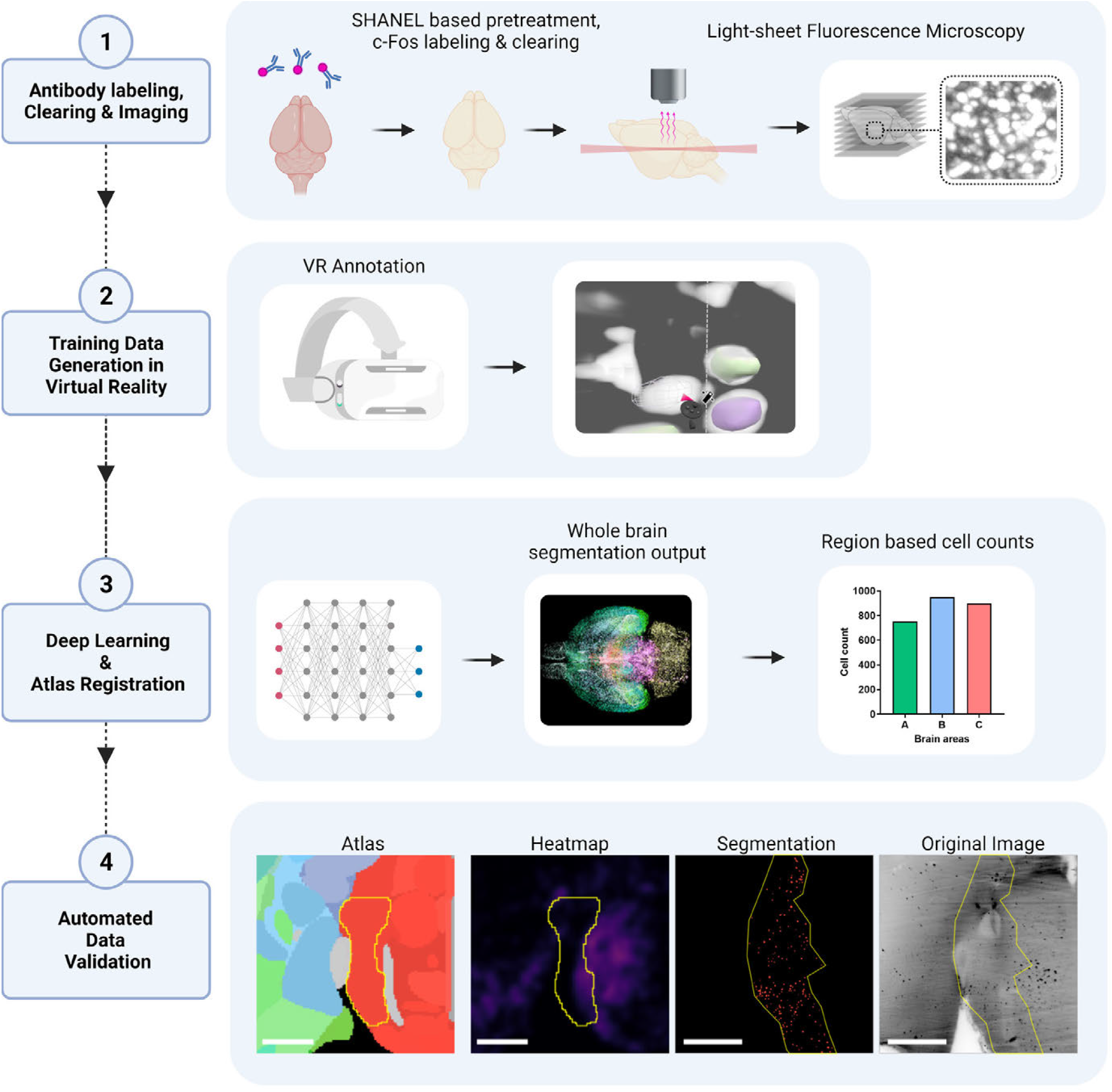
Summary of virtual reality (VR) aided deep learning for antibody labeled cell segmentation in mouse brains. 1. Fixed mouse brains are subjected to SHANEL based antibody labeling, tissue clearing and fluorescent light sheet imaging. 2. Volumes of raw data are labeled in 3D using VR to generate ground truth data. 3. Brains are subjected to deep learning based cell segmentation and registration to the Allen Brain atlas. Subsequently, region based cell counts are extracted and can be analyzed. 4. DELiVR automatically generates a color-coded validation data set.

## RESULTS

### Generation of ground-truth labels is faster and more accurate in VR compared to 2D slice annotation

Deep learning algorithms are constrained by the amount of annotated training data. A more reliable and faster annotation approach can drastically improve deep learning-based image analysis in diverse applications. Here, we aimed to develop a powerful deep learning-based solution to identify c-Fos+ cells. However, such whole-brain immunolabelling datasets present unique challenges, including changes in the ratio of labeling intensity to background intensity in different areas. To train segmentation models in a supervised manner, expert annotations are crucial. Common annotation approaches such as ITK-SNAP10 enable sequential 2D slice-by-slice annotation. Since this is a very time-consuming task, we opted for a VR approach as this allows for full immersion into the 3D volumetric data. We used two different commercial VR annotation software packages (Arivis VisionVR and syGlass^11^), and evaluated their speed and accuracy in comparison to 2D slice annotation in ITK-SNAP on a c-Fos labeled brain sub volume (**Fig. 2a** and **b**). For annotation using Arivis VisionVR, the annotator defined a region of interest (ROI) in which an adaptive thresholding function was applied, according to the annotator’s input. (**Fig. 2c-g, Supplementary Video 1**). In syGlass, the annotation tool allowed the annotator to draw simple three-dimensional shapes as the ROI and adjust a threshold until the annotation was acceptable (**Supplementary Fig. 1a-d, Supplementary Video 2**). In ITK-SNAP, annotation was performed by annotating individual cells in each plane of the image stack (**Supplementary Fig. 1e, Supplementary Video 3**). We evaluated the time spent by the annotators for a 100³ voxel sub volume (depicting 83 cells) as well as the annotation quality of cell instances using the Dice -score. In our experiment, we found that VR annotation was substantially faster compared to 2D slice annotation (**Fig. 2h**) and led to a speed up of annotation time spent of average 7.1 times. Further, VR significantly improved annotation quality, reflected in an increased instance Dice of 80.3% (+6.5%, **Fig. 2i**). In conclusion, VR annotation accelerates label generation compared to 2D slice annotation, leading to more annotated training data of higher quality in in the same time spent. Thus, we decided to generate ground truth data in VR for our deep learning algorithm for c-Fos activity mapping.

**Figure 2:**
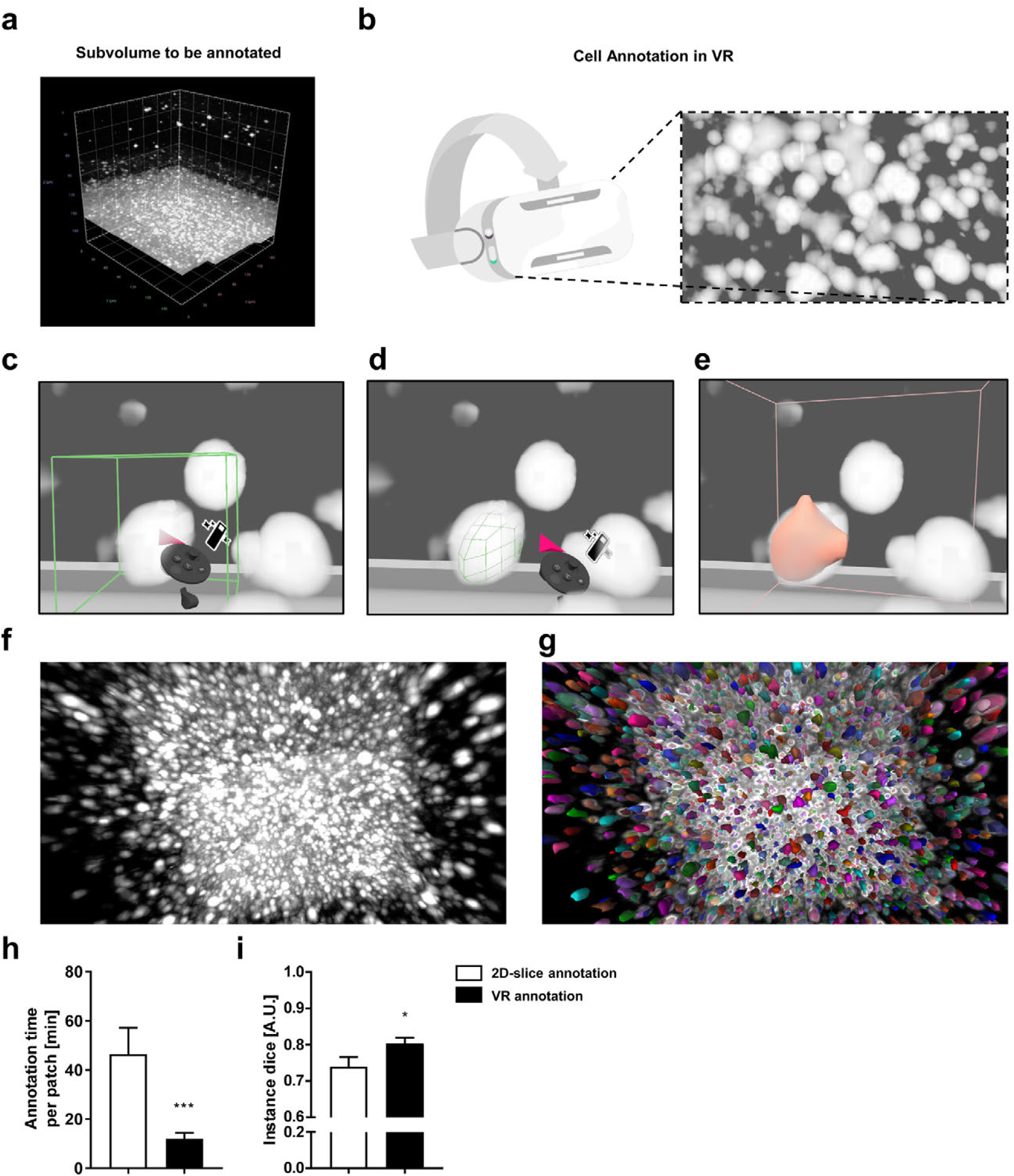
Virtual reality (VR) aided annotation is faster than 2D slice annotation. **a**, Volume of raw data (c-Fos labelled brain) imaged with light-sheet fluorescence microscopy and loaded into Arivis VisionVR. **b**, Illustration of VR goggles and VR view of data. **c-d**, Using Arivis VisionVR, individual cells were annotated by placing a selection cube on the cell (**c**), fitting the cube to the size of the cell (**d**) and filled (**e**). f-g, Zoomed in view of raw data (**f**) and annotation overlay generated in VR (**g**). **h**, Time spent for annotating a test patch on using 2D slice and VR annotation. **i**, Instance Dice of 2D slice annotation vs VR annotation. n=7-12 / group. *p < 0.05, ***p < 0.001

### Deep learning based DELiVR outperforms threshold-based c-Fos segmentation and enables easy visualization of c-Fos expression in brain regions of interest

In order to get a representative sample, we randomly sampled and VR-annotated 48 100³ voxel patches (referring to 5889 cells) from a c-Fos labeled brain that were annotated in VR. From these we randomly selected 9 patches for testing and used 39 patches for training a deep neural network (3D U-Net) with Ranger21 optimizer, MISH activation function and binary cross entropy loss (**Fig. 3a**). Training ran over 500 epochs, with an initial learning rate of 1e-3 and a batch size of 4. We trained the network on a single GPU with 2 random crops of a patch size of 96x96x64 per batch element. We assessed both volumetric as well as instance performance. Volumetric scores are calculated across all voxels in the image, while instance scores are calculated on the overlap between individual cells (see methods section).Our performance on the test set shows a 79.18% instance Dice (+38.66% increase), 84.70% instance sensitivity (+57.79% increase), 65.39% volumetric Dice (+46.56% increase) and 56.85% volumetric sensitivity (+45.39%) compared to ClearMap2 (**Fig. 3b-d, Supplementary Fig. 2a**). These scores demonstrate a clear improvement over filter and threshold-based segmentation methods as the deep learning model captures 73 times more cells (1611 true positives) than ClearMap1 (22 true positives) and 3 times more cells than ClearMap2 (515 true positives) while not oversegmenting (**Fig. 3e**).

**Figure 3:**
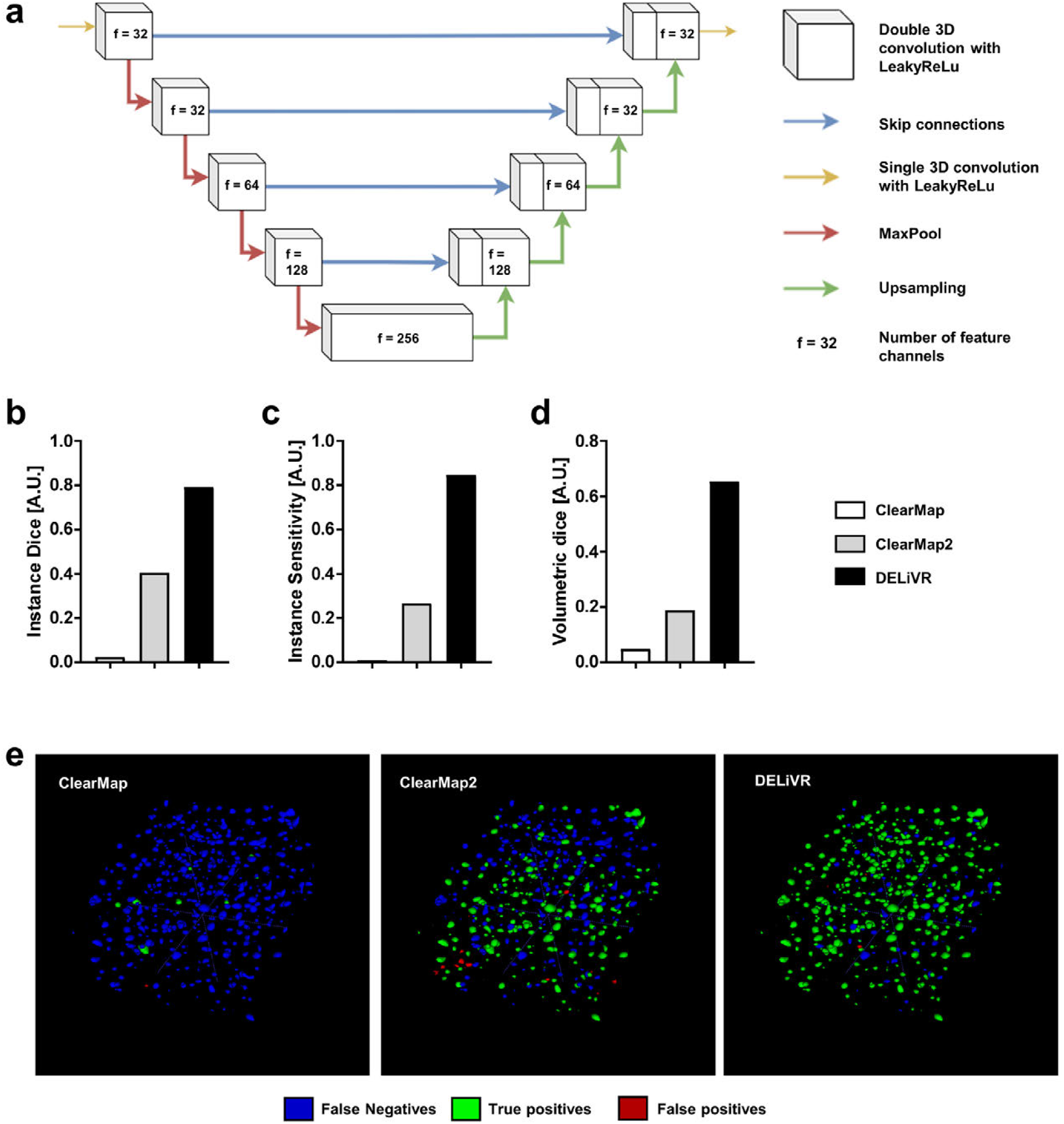
Deep learning-based DELiVR outperforms current methods for c-Fos detection. **a**, Architecture of the c-Fos deep learning network: a MONAI BasicUNet3D. **b – d**, Quantitative comparison between ClearMap, ClearMap2 and our model based on instances of cells and volumetric dice. **e**, 3D qualitative comparison between ClearMap, ClearMap2, and our model on instance basis: Predicted cells with overlap in ground truth are masked in green, predicted cells with no overlap in ground truth are masked in red, and ground truth cells with no corresponding prediction are marked in blue.

Whole-brain antibody labeling with c-Fos often leads to antibody accumulations in ventricles of the brain, thereby generating artifacts in these areas. To exclude these areas from the analysis, we developed an automated pre-processing step that masks the ventricles (**Supplementary Fig. 2b-e**). DELiVR then uses a customized sliding window inferrer for the forward pass. Afterwards we conduct a connected component analysis^12^ to identify individual cells. After atlas alignment and size filtering of the cells, we assign the corresponding atlas region to each found cell and for visualization dilate the cells with a kernel of size 4. The connected component analysis returns a set of center-point coordinates and volume for each segmented cell, which DELiVR then automatically maps to the Allen Brain Atlas (CCF3, 50 µm/voxel) with mBrainAligner^13^. Thereby, DELiVR generates a whole-brain segmentation output that exists in the original image space. Here, each segmented cell corresponds to a threshold value fitting to an Area ID of the Allen Brain Atlas and was colored according to the brain region it belongs to (**Fig. 4a and b, Supplementary Video 4**). In addition, we used BrainRender^14^ to plot and visualize the detected cells in the atlas space (**Fig. 4c**).

**Figure 4:**
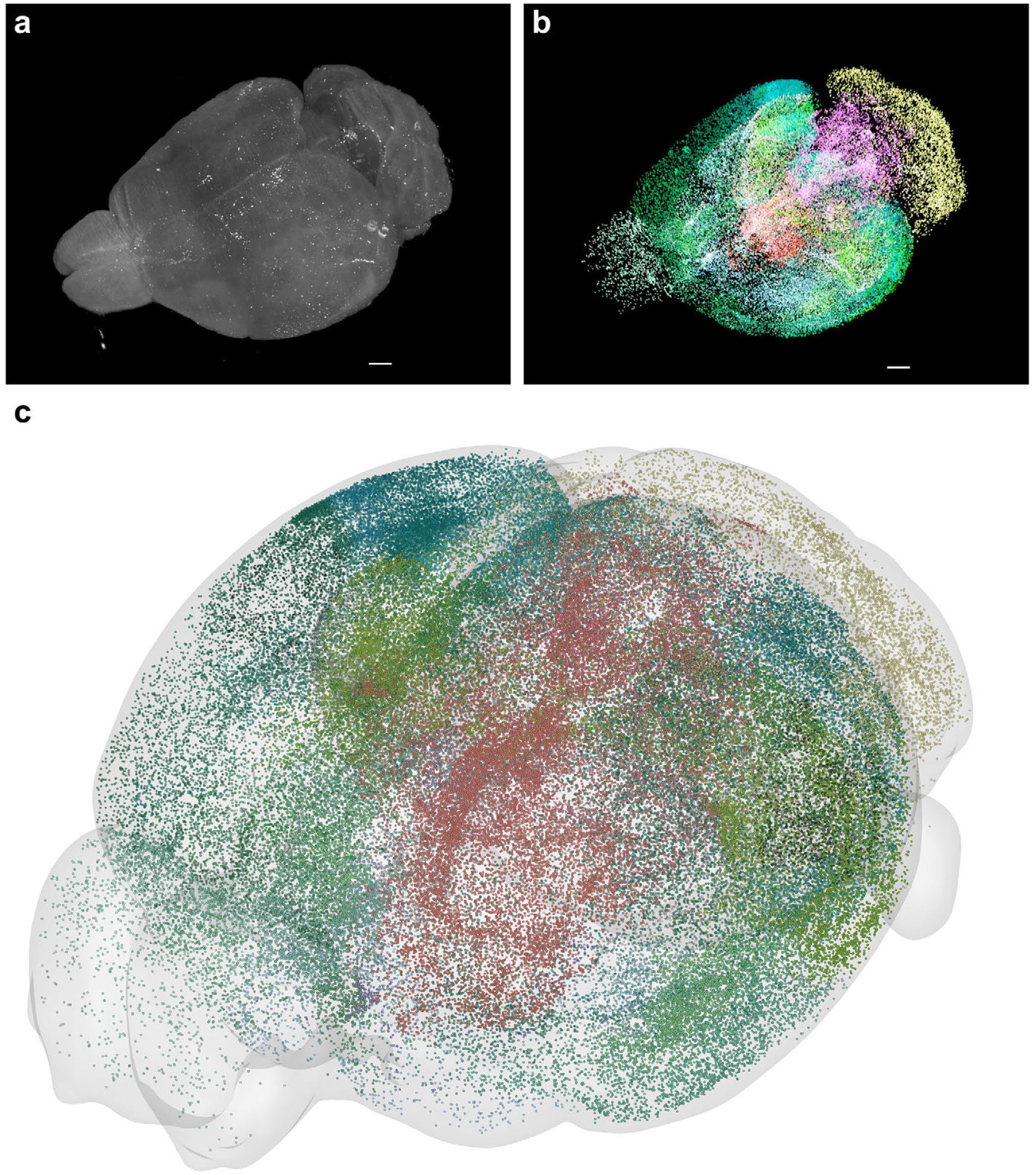
Whole-brain segmentation output generated with DELiVR. **a-b**, 3D view of whole brain (**a**) and segmentation of detected cells, color-coded by atlas region (**b**). **c**, visualization of the area-mapped cells in Atlas Space with BrainRender.

### DELiVR identifies a novel brain activation pattern in weight-stable tumor-bearing mice

Cancer affects the normal physiology of cells leading to changes in the metabolic activity of the tumor and surrounding tissues. It can also lead to profound changes in systemic metabolism of the patient. This is particularly exemplified in the wasting syndrome cancer-associated cachexia (CAC) that is characterized by involuntary loss of body weight^15-17^ and specific changes in brain activity^18^. To identify brain regions that are potentially involved in affecting body weight maintenance in cancer we used DELiVR to compare the neuronal activity patterns between weight-stable cancer and CAC. We subcutaneously transplanted NC26 colon cancer cells that give rise to weight-stable cancer or C26 colon cancer cells, which induce weight loss (**Fig. 5a**). As expected, no changes in body weight were observed in NC26 tumor-bearing mice compared to controls, while C26 tumor-bearing mice showed significant reductions in body weight (**Fig. 5b**). The differences in body weight were not due to differences in tumor mass, as the tumor size was similar between NC26 and C26 tumor-bearing animals (**Fig. 5c**). C26 tumor-bearing mice also displayed reduced weights of the gastrocnemius muscle and white adipose tissue depots (**Supplementary Fig. 3a-c**). We performed c-Fos antibody labeling, clearing and imaging of whole brains of these mice and applied DELiVR for unbiased whole-brain mapping of neuronal activity. Interestingly, c-Fos+ density maps indicated an increase in brain activity in weight-stable NC26 tumor-bearing mice compared to PBS controls, while this increase was not present in cachectic C26 tumor-bearing mice (**Fig. 5d**).

**Figure 5:**
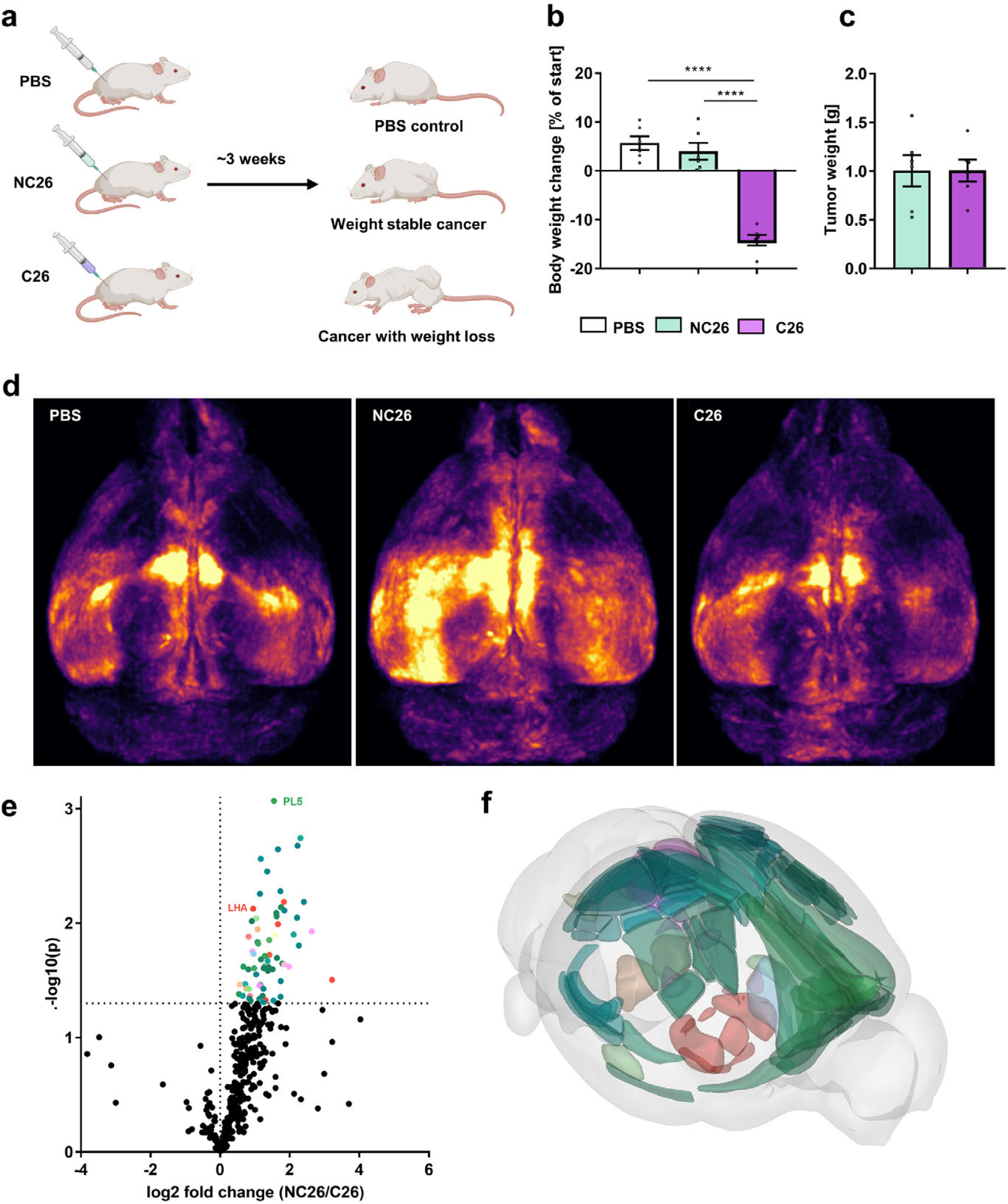
DELiVR identifies changes in neuronal activity in weight-stable cancer. **a**, Experimental set-up. Mice were subcutaneously injected with PBS, NC26 cells that lead to a weight-stable tumor formation or cachexia-inducing C26 cancer cells. **b**, Mouse body weight change over the course of the experiment. **c**, Tumor weight at the end of the experiment. **d**,, Average density maps of c-Fos expression in brains of PBS controls, mice with weight-stable cancer (NC26) and mice with cancer-associated weight loss (C26). **e**, Volcano plot of areas activated in weight-stable cancer (NC26 over C26) based on c-Fos expression. Significantly changed areas are color-coded according to the Allen Brain Atlas. **f**, Significantly different areas between NC26 and C26, visualized using BrainRender. n=6/group. ****p < 0.0001.

Further, we found significantly increased c-Fos+ density in 73 areas in the NC26 brain compared to C26 (**Fig. 5e-f, Fig. 6a)**. We found approximately half of the overactive cortical areas in NC26 tumor-bearing mice are linked to higher sensory processing, such as primary and secondary visual, auditory and somatosensory, areas as well as the ectorhinal cortex (**Fig. 6a**). In addition, regions associated with foraging such as the retrosplenial and prelimbic cortex were hyper-activated in NC26 tumor-bearing mice (**Fig. 6a)**. Besides cortical areas, we observed increased c-Fos expression in several hypothalamic nuclei including the lateral hypothalamic nucleus (LHA), which is an important regulator of feeding and metabolism^19,20^. To confirm the validity of our quantifications, we used DELiVR’s novel visualization tool for c-Fos expression confirmation in brain regions of interest. To this end, we evaluated c-Fos expression in our original images and in the DELiVR segmentation output in the LHA and prelimbic areas in brains of weight-stable and cachectic tumor-bearing mice (**Fig 6b**). The color mapping of the cells allows to highlight the area of interest e.g., using standard Fiji tools. Doing so, we were able to highlight only the segmented cells of this region and thereby making it easy to find and confirm an anatomical or functional sub-area in the original image stack of the brain. In agreement with the quantification in Fig 6a, we observed increased c-Fos expression in the LHA and prelimbic area of NC26 tumor-bearing mice (**Fig. 6b**).

**Figure 6:**
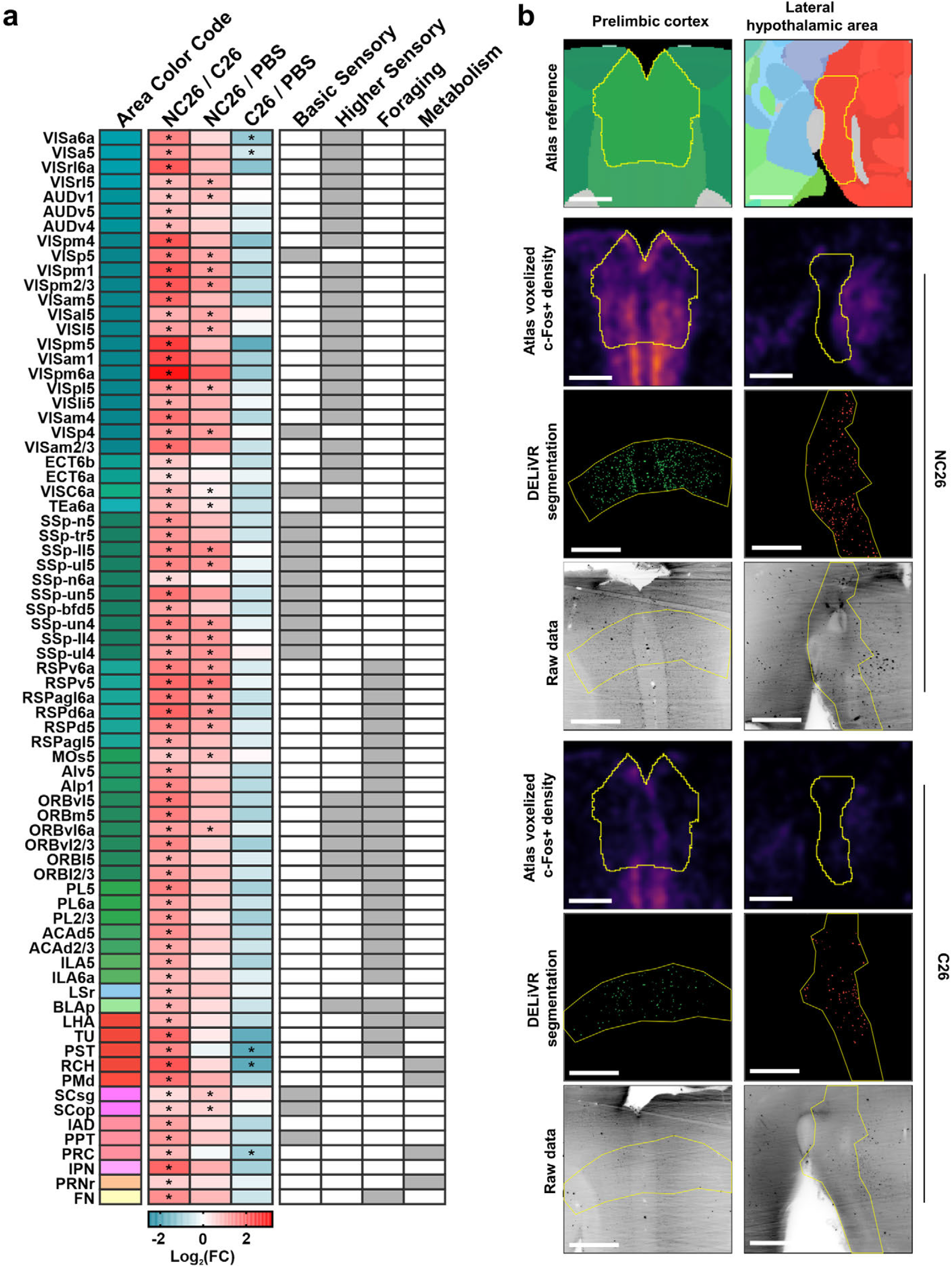
DELiVR identifies cancer-related brain activation patterns. **a**, Fold change of brain regions induced by cancer compared to a control group. * p<0.05 (n=6/group). **b**, Per-region cell label validation in the prelimbic Cortex (PL) and Lateral Hypothalamic Area (LHA). Atlas and average heatmap are displayed in atlas space (25µm/px). The labelled cells and microscope images are displayed in the original image Space (36 µm z-projection). Scale bars = 500 µm in both spaces.

Overall, our findings showed that brain activity in weight-stable NC26 cancer-bearing mice are markedly different from both cachectic C26 cancer-bearing mice and PBS controls. Thus, with DELiVR we were able to identify a novel neuronal activity pattern specific to the NC26 cancer model.

## DISCUSSION

DELiVR is a VR-enabled deep learning-based quantification pipeline to study brain activity in cleared mouse brains. We used VR to generate ground truth data for training a deep learning based segmentation network. DELiVR improves segmentation accuracy compared to current cell detection methods and generates a registered segmentation output that can be examined in the original image and in the atlas spaces.

Our experiments show that VR is more than a medium for immersive games, but rather a superior means of annotation and data exploration for volumetric data analysis. Non-VR methods for segmentation show orthogonal slices, which allows an annotator to outline the shape of individual cells in 2D. However, it obfuscates necessary spatial information which makes annotation challenging and time consuming - an annotator never sees the cell as a whole, only a cross section, and has to scroll through slices to ensure that it is in fact a cell and not background noise. In contrast, VR allows the annotator to capture three-dimensional structures in their entirety and thereby enables the fast generation of more reliable annotated data.

Traditional, non-machine learning computer vision solutions for large-scale analysis of c-Fos cell detection such as ClearMap^6,21^ rely on a sophisticated system of thresholding and filtering to detect small structures and classify them as cells. While such approaches have been proven to generate valuable information5, they show limited performance for data with variable signal-to-noise ratios, as is the case when imaging large volumes such as the mouse brain using LSFM. Those parameters can be adjusted, but it is very difficult to find a parameter set which accounts for all cells and hence the thresholds tend to be set conservatively, meaning that subtle differences may be lost during threshold-based analysis. A trained deep learning model however learns these local variances, allowing for a one-shoe-fits-all solution once trained sufficiently. We find that with our datasets, DELiVR increases instance Dice, volumetric Dice as well as instance sensitivity. Thus, DELiVR can fulfill the desired task of giving us the correct number of cells in the right locations, while threshold-based cell detection missed a large number of cells by being too conservative.

We used DELiVR to profile the brain activation patterns of cancer-bearing mice that were either weight-stable or displayed CAC. Weight loss in cancer is driven by a mix of reduced food intake, elevated catabolism, increased energy expenditure and inflammation^17^. The brain was shown to contribute to anorexia in CAC, as it responds to inflammatory cytokines that modulate the activity of neuronal populations that regulate appetite^22^. In addition, activation of neurons in the parabrachial nucleus was shown to suppress appetite in mouse models of CAC^23^. Of note, decreased food intake alone is not the sole reason for cachexia development as nutritional support fails to fully revert the syndrome^24^. In our dataset, we found increased c-Fos+ cell counts in the LHA of weight-stable NC26 tumor-bearing mice compared to cachectic C26 tumor-bearing mice, indicating a connection between LHA activity and weight loss. Specifically in cachexia, reduced LHA activity is caused by hypothalamic inflammation, which triggers silencing of orexinergic neurons, thereby tilting the energy balance towards energy wasting ^18,25,26^. In contrast, the NC26-specific LHA hyperactivation could indicate a novel compensatory mechanism that maintains normal body weight in this cancer model.

Surprisingly, we also found a strong increase in c-Fos+ counts in brains of weight-stable NC26 tumor-bearing mice in numerous other locations. Many of these areas where located in the cortex and included all primary sensory regions (somatosensory, visual, and auditory), as well as secondary sensory regions, the retrosplenial cortex, and large parts of the prelimbic and orbitofrontal regions – all involved in higher-order sensory processing as well as motion sequencing and (partially) foraging^27-29^. The abundance of sensory-related regions being affected in our weight-stable cancer model also opens the possibility of a cancer-specific impairment in GABAergic inhibition^30^. The strong sensory component is unique to NC26 non-cachectic cancer brains in our dataset and differentiates them from both C26 and PBS controls. Whether this increase in neuronal activation in weight-stable cancer bearing mice somehow affects body weight maintenance will be interesting to explore in future studies.

To conclude, we present DELiVR: an integrated pipeline to label, scan, and analyze neuronal activity markers across the entire mouse brain. We innovate by bringing a deep-learning segmenter to cleared-brain analysis, and speed up processing time by generating the required training data more accurately in VR. Using DELiVR, we find differences in c-Fos expression between cachectic and non-cachectic cancer mouse brains, pointing us to a previously unknown neurophysiological phenotype in cancer-related weight control.

## MATERIALS AND METHODS

### Whole-brain immunolabeling and clearing

Immunostaining for c-Fos was performed using a modified version of SHANEL9. All incubation steps were carried out under moderate shaking (300 rpm). For the pretreatment, samples were dehydrated with an ethanol/water series (50% EtOH, 70% EtOH, 100% EtOH) at room temperature for 3h per step. Next, samples were incubated in DCM/MeOH (2:1 v/v) at room temperature for 1 day. Brains were rehydrated with an ethanol/water series (100% EtOH, 70% EtOH, 50% EtOH, diH20) at room temperature for 3h per step. Samples were incubated in 0.5M acetic acid at room temperature for 5 hours followed by washing with diH_2_O. Next, brains were incubated in 4M guanidine HCl, 0.05M sodium acetate, 2% v/v triton x100, pH=6.0 at RT for 5 hours followed by washing with diH_2_O. Brains were incubated in a mix of 10% CHAPS and 25% N-Methyldiethanolamine at 37° for 12h before washing with diH_2_O. Blocking was performed by incubating the brains in 0.2% triton x100, 10% DMSO, 10% goat serum in PBS shaking at 37° for 2 days. Samples were incubated with c-Fos primary antibody (Cell Signaling, #2250, 1:1000) in primary antibody buffer (0.2% Tween-20, 5% DMSO, 3% goat serum, 100ul heparin/100 ml PBS) shaking at 37° for 7 days. The antibody solution was filtered (22 µm pore size) before use. Samples were washed in washing solution (0.2% Tween-20, 100ul heparin/100 ml PBS) shaking at 37° for 1 day at which the washing solution was refreshed 5 times. Brains were incubated with the secondary antibody (Alexa Fluor 647, Goat anti-Rabbit IgG (H+L) from Invitrogen #A-21245) in secondary antibody buffer (0.2% Tween-20, 3% goat serum, 100ul heparin/100 ml PBS) shaking at 37°C for 7 days followed by incubating in washing solution shaking at 37° for 1 day at which the washing solution was refreshed 5 times.

Brains were dehydrated using 3DISCO2 with a THF/diH_2_0 series (50% THF, 70% THF, 90%THF, 100%THF) for 12h per step followed by an incubation in DCM for 1h. Tissues were incubated in benzyl alcohol/benzyl benzoate (BABB, 1:2 v/v) until tissue transparency was reached (>4 h).

### Light-Sheet Imaging

Light-sheet imaging was conducted through a 4× objective lens (Olympus XLFLUOR 340) equipped with an immersion corrected dipping cap mounted on an UltraMicroscope II (LaVision BioTec) coupled to a white light laser module (NKT SuperK Extreme EXW-12). The antibody signal was visualized using a 640/40 nm excitation and 690/50 nm emission filter. Tiling scans (3 by 3 tiles) were acquired with a 20% overlap, 60% sheet width and 0.027 NA. The images were taken in 16 bit depth and at a nominal resolution of 1.625 μm/voxel on the XY axes. In z-dimension we took images in 6 μm steps using left and right sided illumination. Stitching of tile scans was carried out using Fiji’s stitching plugin, using the “Stitch Sequence of Grids of Images” plugin31 and custom Python scripts.

### ClearMap

ClearMap6 and ClearMap2’s CellMap portion^21^ were used with adapted settings for thresholds and cell sizes that fitted to the higher resolution and different signal to noise ratios in our dataset. Segmentation masks were saved as tiff stacks by toggling the “save” option in the last segmentation step. ClearMap 1 was ported to Python 3 before use, but functioned identically32. We only used the cell segmentation portions, no pre-processing (e.g. ClearMap2’s flat-field correction) or post-processing such as atlas alignment were performed. Both pipelines were run for an entire brain and subsequently subdivided into test patches that we used for the comparisons with DELiVR.

### Ventricle masking

We wrote an automated preprocessing script that downsamples the image stack to an isotropic 25x25x25 µm/voxel, and then applies a custom-trained Random Forest to identify ventricles. Specifically, we integrated Ilastik^33^ (version 1.407b) with a 3D pixel classifier, which we trained on several downsampled brain image stacks to differentiate between ventricles and brain parenchyma. The preprocessing script then generates a 3D mask stack, which our script upsamples to the original image stack dimensions, using bicubic interpolation to avoid aliasing artefacts at ventricle edges. It then masks each original z-plane image with the respective mask and returns a 16-bit tiff stack with the ventricles masked out.

### Annotation

VR annotation was carried out using Arivis VisionVR or syGlass^11^. For this purpose, the annotator was wearing a VR head set (Oculus Rift S) and carried out annotations in VR using hand controllers (Oculus touch). Slice by slice annotation was carried out using ITK-SNAP^10^. For comparing VR and 2D-sliced based annotation, a 1003 voxel volume of c-Fos labelled brain was annotated by the participants and the time was recorded until the annotation task was finished. For training and testing our deep learning network, we annotated a total of 48 100³ voxel patches in VR. All of our training and test patches were furthermore vetted by an expert biologist in ITK-SNAP to ensure that only cells were annotated. We evaluated the annotation quality using the formula of Dice as described below.

### Deep Learning

To automatically segment the cells in all brains we trained a BasicUnet3D from the MONAI library^34^. The annotated dataset of 48 100³ patches was split in 9 patches for testing and 39 patches for training. As an activation function we chose MISH35 and as optimizer Ranger2136 in order to flatten the loss landscape and allow for easier generalization. As loss function we used binary cross entropy loss. For the training of 500 epochs, we set the initial learning rate to 1e-3 and the batch size to 4. The network is then trained on a single GPU (Nvidia RTX3090) with 2 random crops with a patch size of 96x96x64 per batch element.

### Evaluation of the segmentation model

Evaluation of the deep learning model was done in a two-fold way, we compared both the volumetric quality of the segmentation by assessing for each voxel if it was correctly classified as foreground or background, as well as the instance segmentation quality, where we assess on a single cell level whether a cell has been detected or not. Volumetric quality assessment gave us true positives (TP), false positives (FP), false negatives (FN) and true negatives (TN) by comparing every prediction voxel with the ground truth voxel. Instance quality assessment gave us TP, FP and FN by first conducting a connected component analysis on both the network prediction as well as the ground truth. For each connected component, we then compared whether there was at least a single voxel overlap between one connected component in the prediction and in the ground truth. The corresponding ground truth component was then removed from the set of components and was counted as true positive. Every ground truth component that had no overlap was counted as a FN, every prediction component without overlap was counted as a FP. The intuition here is that our biological question requires a truthful count of cells at the correct area rather than voxel perfect segmentations of each cell. Comparison with ClearMap and ClearMap2 was performed as follows: We first applied the methods on the same brain we took our test data from, masked it with our random forest masker and then cut out the relevant 100³ voxel patches. These patches were then compared to our ground truth annotation.

Scores were defined as follows:

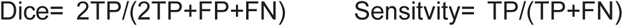

### Atlas registration and statistical analysis

Because the SHANEL protocol’s aggressive detergent treatment warps the brains considerably more than in other clearing techniques, we could not use Elastix for Atlas registration (data not shown). Rather, we used a novel and recently published tool, mBrainAligner^13^, which worked well with our datasets (Supplementary Fig. 4). We manually saved the downsampled isotropic 25x25x25 µm/voxel stacks as .v3draw using Vaa3d^37^. Subsequently, we wrote an automated script that aligned the image stacks to mBrainAligner’s 50x50x50 µm/voxel version of the Allen Brain Atlas CCF3 reference atlas, using the Light-Sheet Fluorescent Microscopy example settings with minor adaptations. Subsequently, we used mBrainAligner’s swc transformation tool to map the center point coordinates of our c-Fos+ cells into atlas space.

Furthermore, we wrote a custom cell-to-atlas script (reusing parser code from VeSSAP^38^, and the CCF3 atlas file as provided by the Scalable Brain Atlas^39^) that filters the cells by size, based on 3x standard deviation (upper bound = 104 voxels) and returns two tables: A table with each cell as a row, including the region, Allen Brain Atlas color code, etc. Furthermore, a region table with one region per row, in which the number of c-Fos+ cells per region is summarized. Lastly, for all datasets, the postprocessing script generates overview tables that contain cell counts for all regions. We used the latter for uncorrected Student’s t tests. Lastly, we implemented a level-aware multiple-testing script that compares between groups at the Allen Brain Atlas’s 11 structure levels.

### Visualization

For visualizing the cells and regions in atlas space, we used BrainRender^14^ with a modified density plot function^32^. In order to visualize the segmented cells in the original image space, we combined the area-wise color-code from the Allen Brain Atlas with the 3D segment mask output by the connected component analysis. The result is a cell mask file with each cell being color-coded according to the brain area it belongs to, which makes overlaying with the original image data in e.g. Fiji easy and allows for direct visual inspection of the segmentation results.

### Cell culture

C26 and NC26 colon cancer cells were cultured in high glucose DMEM with pyruvate (Life Technologies #41966052), supplemented with 10% fetal bovine serum (Sigma-Aldrich #F7524) and 1% penicillin-streptomycin (Thermo Fisher #15140122)^40^. Before using the cells for transplantation, cells had a confluence of 80%. Cells were trypsinized, counted and required cell numbers were suspended in Dulbecco’s phosphate-buffered saline (PBS, Thermo Fisher #14190250).

### Animal experimentation

Experiments were carried out with male BALB/c mice at an age of 10-12 weeks. They were purchased from Charles River (CRL, Brussels), maintained on a 12-h light–dark cycle and fed a regular unrestricted chow diet. They were injected with 1x106 C26 or 1.5x106 NC26 cells in 50 µl PBS subcutaneously into the right flank. Control mice were injected with PBS. 5 days after cell implantation, mice were monitored daily for tumor growth and body weight. Cachectic C26 tumor-bearing mice considered cachectic when they had lost 10% - 15% of body weight. Mice were sacrificed following deep anesthesia with a mix of ketamine/xylazine, followed by intracardiac perfusion with heparinized PBS (10 U/ml heparin) followed by a perfusion with 4% paraformaldehyde (PFA). Tissues and organs were dissected, weighed and post-fixed at 4°C overnight. Animal experimentation was performed in accordance with the European Union directives and the German animal welfare act (Tierschutzgesetz). They have been approved the state ethics committee and the government of Upper Bavaria (ROB-55.2-2532.Vet_02-18-93).

### Statistical analysis

Results from biological replicates were expressed as mean ± standard error of the mean. Statistical analysis was performed using GraphPad Prism 7. Normality was tested using Shapiro–Wilk normality tests. To compare two conditions, unpaired Student’s t-tests or Mann– Whitney tests were performed. One-way analysis of variance (ANOVA) or Kruskal–Wallis tests with Sidak’s post hoc test were used to compare three groups.

## Supporting information

Supplementary Video 1. Annotation of cells in virtual reality using Arivis Vision VR

Supplementary Video 2. Annotation of cells in virtual reality using syGlass

Supplementary Video 3. 2D-slice based annotation using ITK-snap

Supplementary Video 4. Whole brain with c-Fos+ cells detected by DELiVR depicted in the original image space. Cells are color coded by area they belon

## Acknowledgements

We would like to thank AIME (Berlin) for processing time on their servers. We thank Izabela Horvath, Hongcheng Mai and Louis Kümmerle from the Institute for Tissue Engineering and Regenerative Medicine (Helmholtz Munich) for annotation tasks. We thank Luke Harrison from the Institute for Diabetes and Cancer (Helmholtz Munich) for editing the graphical summary. Fig.1 and Fig.5a were generated using Biorender. We thank Rudolf Zechner and Martina Schweiger from the Institute of Molecular Biosciences (University of Graz) for kindly providing NC26 cancer cell lines. This work was supported by a grant from the Else-Kröner-Fresenius-Stiftung (2020 EKSE.23) to S.H..

## Author contributions

DK, RA, SH, MBD and AE conceptualized the study. DK performed in vivo work, antibody labeling, imaging and data processing. RA performed VR and deep learning analysis. MN performed atlas registration, data analysis and interpreted the results. LH performed VR annotations. FK supported the deep learning analysis. SZ and MT supported imaging and antibody labeling. JG and PM supported in vivo experiments. JCP supported data interpretation. MR provided funding and useful discussions. BW provided funding. SH supported supervision of the study. AE and MBD supervised the study. DK, RA und MN drafted the manuscript. AE, DK, MN and RA revised the manuscript.

## Data and code availability

Our training and test data as well as the trained network and all code to run our pipeline end-to-end is available under https://github.com/erturklab/delivr_c-Fos/. A Docker image of the complete pipeline will be available in the near future. Raw data are available on reasonable request.

---

**Suppl. Fig. 1.**
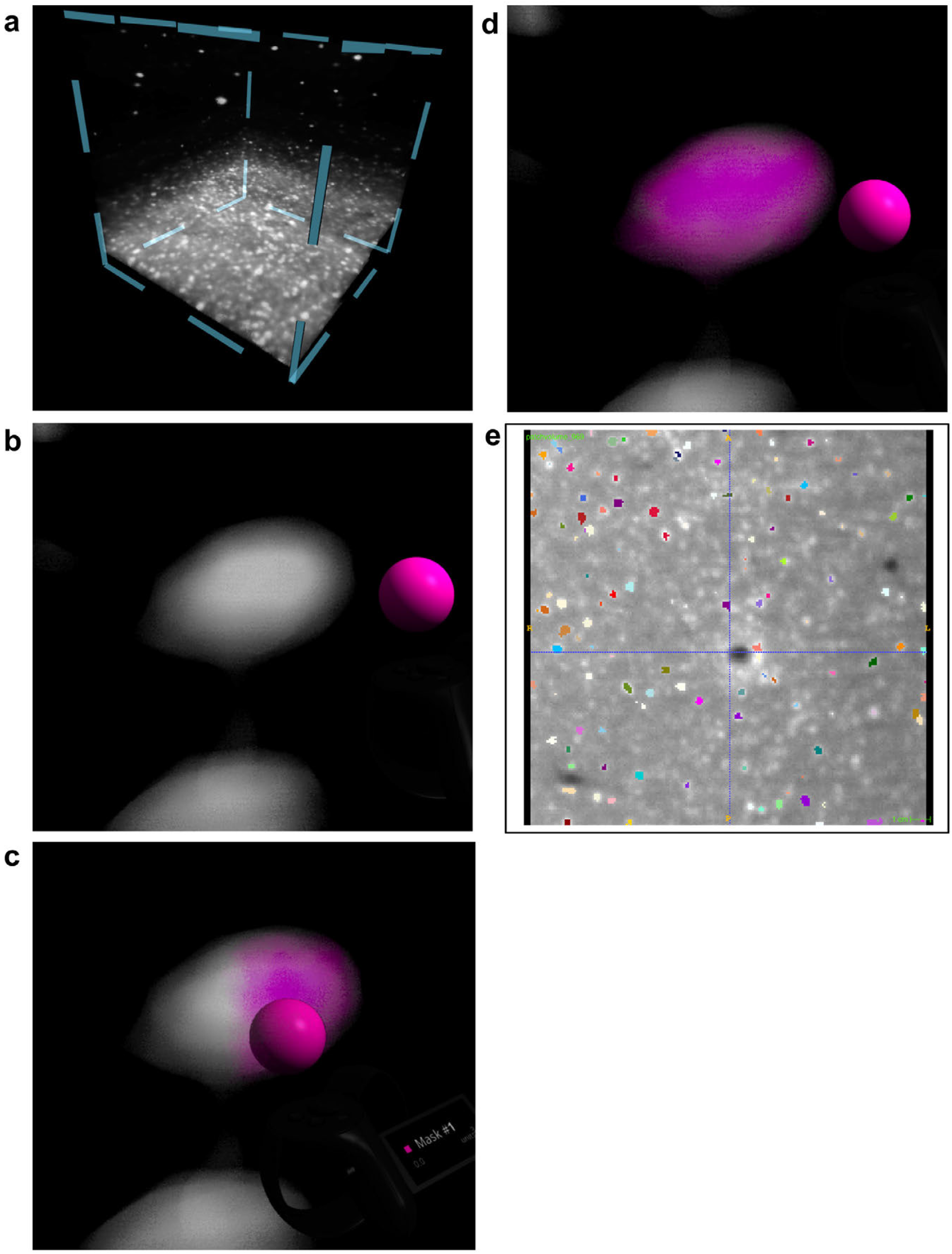
VR Segmentation in syGlass and 2D slice-based segmentation using ITK-SNAP. **a**, Volume of raw data (c-Fos labelled brain) that was generated by light sheet microscopy and loaded into syGlass. **b-d**, Using VR, individual cells were segmented in syGlass by using three-dimensional euclidean shapes as ROI and adjust a threshold until the segmentation was acceptable. **e**, ITK-SNAP view of a single plane of the image stack. Cells were labelled individually slice by slice. Colored cells indicate the segmentations performed by the annotator.

**Suppl. Fig. 2.**
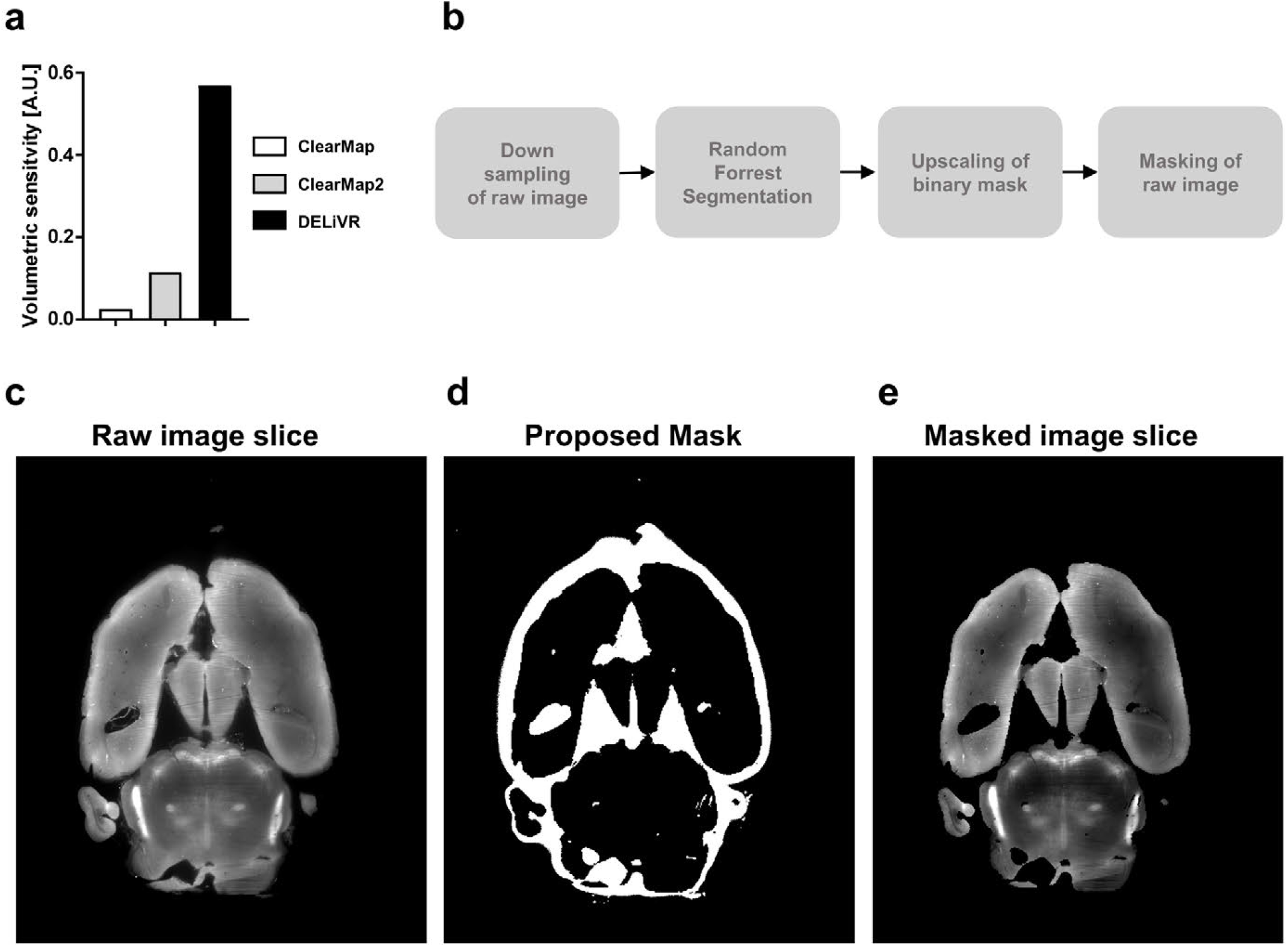
DELiVR pre-processing automatically removes artefacts. **a**, Quantitative comparison between ClearMap, ClearMap2 and DELiVR based on volumetric sensitivity. **b**, Preprocessing workflow. DELiVR preprocessing down-samples the original image stack. Subsequently, it uses a random forest segmenter (Ilastik pixel classification) to create a mask for ventricles. Lastly, the mask is upscaled and applied on the original slices to create a masked image stack. **c-e**, Horizontal view of an original image slice (**c**), the proposed mask (**d**) and the masked image slice generated (**e**).

**Suppl. Fig. 3.**
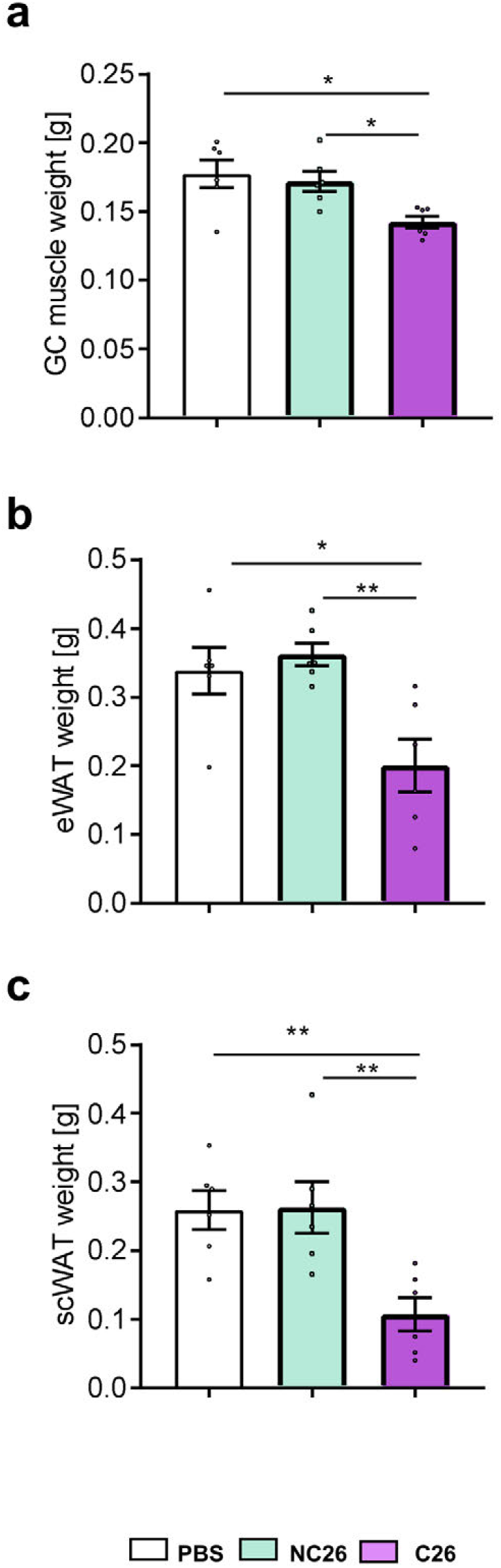
Tissue weights of mice with weight-stable cancer (NC26) and cancer-associated weight loss (C26) **a-c**, Weights of gastrocnemius (GC) muscle (**a**), epididymal white adipose tissue (WAT) (**b**) and subcutaneous WAT (**c**). n=6/group. *p < 0.05, **p < 0.01.

**Suppl. Fig. 4.**
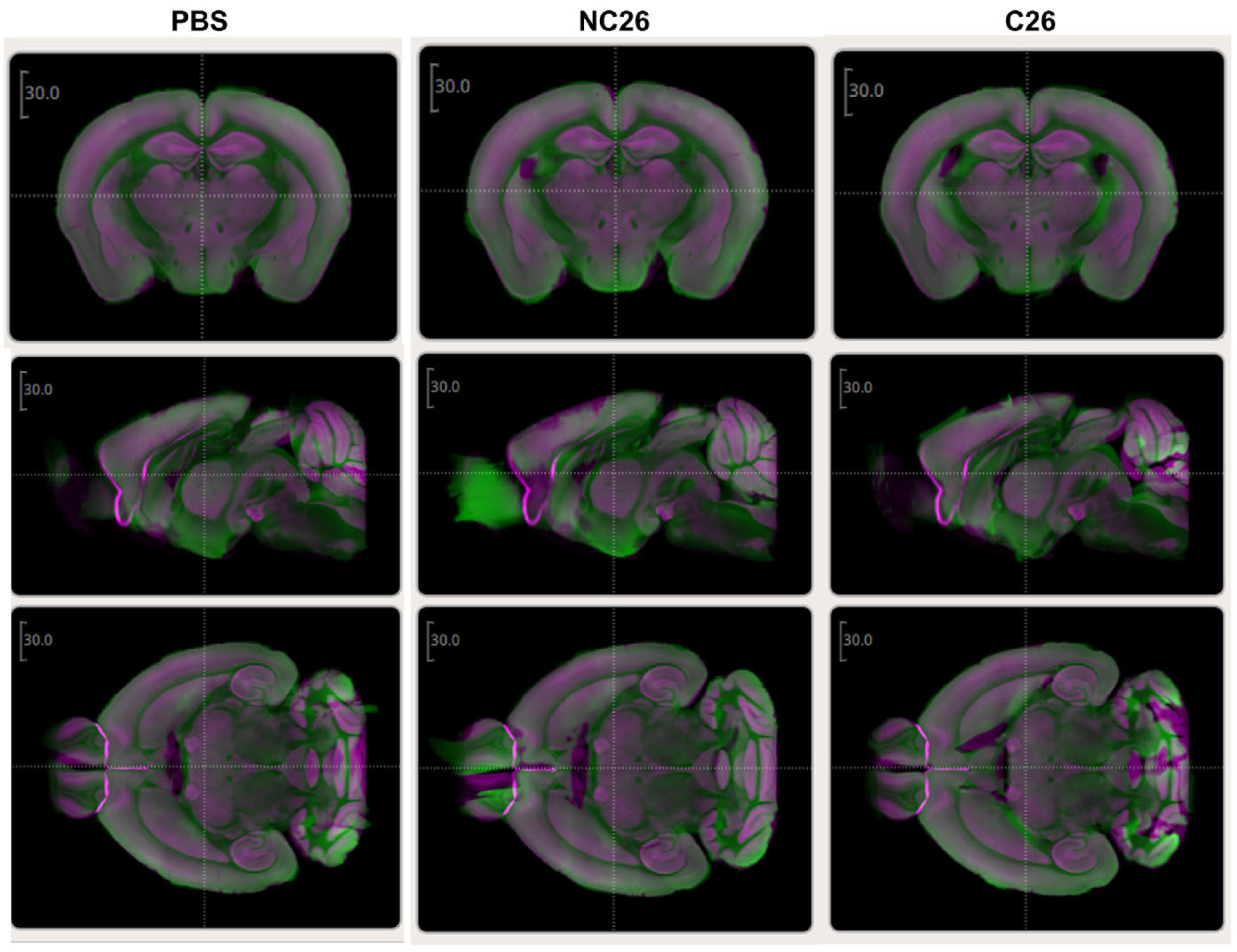
Atlas Registration with mBrainAligner. Post-registration overlap for representative brains of the experimental groups PBS (left), NC26 (middle) and C26 (right). Green, registered image stack. Magenta, mBrainAligner’s version of the CCF3 reference atlas.

